# Stage-adaptive integration of polydopamine promotes human pluripotent stem cell-derived alveolar organoids differentiation and maturation

**DOI:** 10.64898/2026.03.02.708928

**Authors:** Ruihao Lan, Yu Chen, Zhiying Liao, Hengrui Zhang, Caidie Zhong, Jiaxiang Yin, Chang Du, Tao Xu, Hao Meng, Huisheng Liu

## Abstract

Human pluripotent stem cell (hPSC)-derived alveolar organoids (ALOs) have emerged as a powerful tool for modeling human lung development and disease, and accelerating respiratory drug discovery. However, achieving the functional maturation of ALOs remains challenging. Polydopamine (PDA) is a mussel-inspired polyphenolic biomaterial with antioxidant and adhesive properties that can be deployed as surface coatings and nanoparticles (NPs) in cell culture systems. Here, we integrate PDA coatings and NPs sequentially in a stage-adaptive manner throughout the hPSC-derived ALOs differentiation system and study their contributions to ALOs maturation. Our results demonstrated PDA coating yielded more anterior foregut endoderm (AFE) spheroids by strengthening the interaction between Matrigel and substrate. Bulk RNA-seq revealed enrichment of cell–cell and cell-extracellular matrix interactions by PDA. The subsequent incorporation of PDA NPs in Matrigel at lung progenitor cells (LPCs) stage significantly mitigated reactive oxygen species (ROS) accumulation and enhanced LPCs generation. Functionally, AT2 cells in ALOs exhibit characteristic lysosome-to-lamellar body (LB) maturation due to the traffic of internalized PDA NPs to endolysosome. Transcriptomics further indicated enrichment of endocytic-phagosome and epithelium development pathways by PDA treatment. Together, our study establishes a stage-adaptive-integrated PDA strategy throughout hPSC-to-ALOs differentiation and demonstrates that PDA robustly enhances ALOs maturation and secretory function.

**Graphic abstract:** 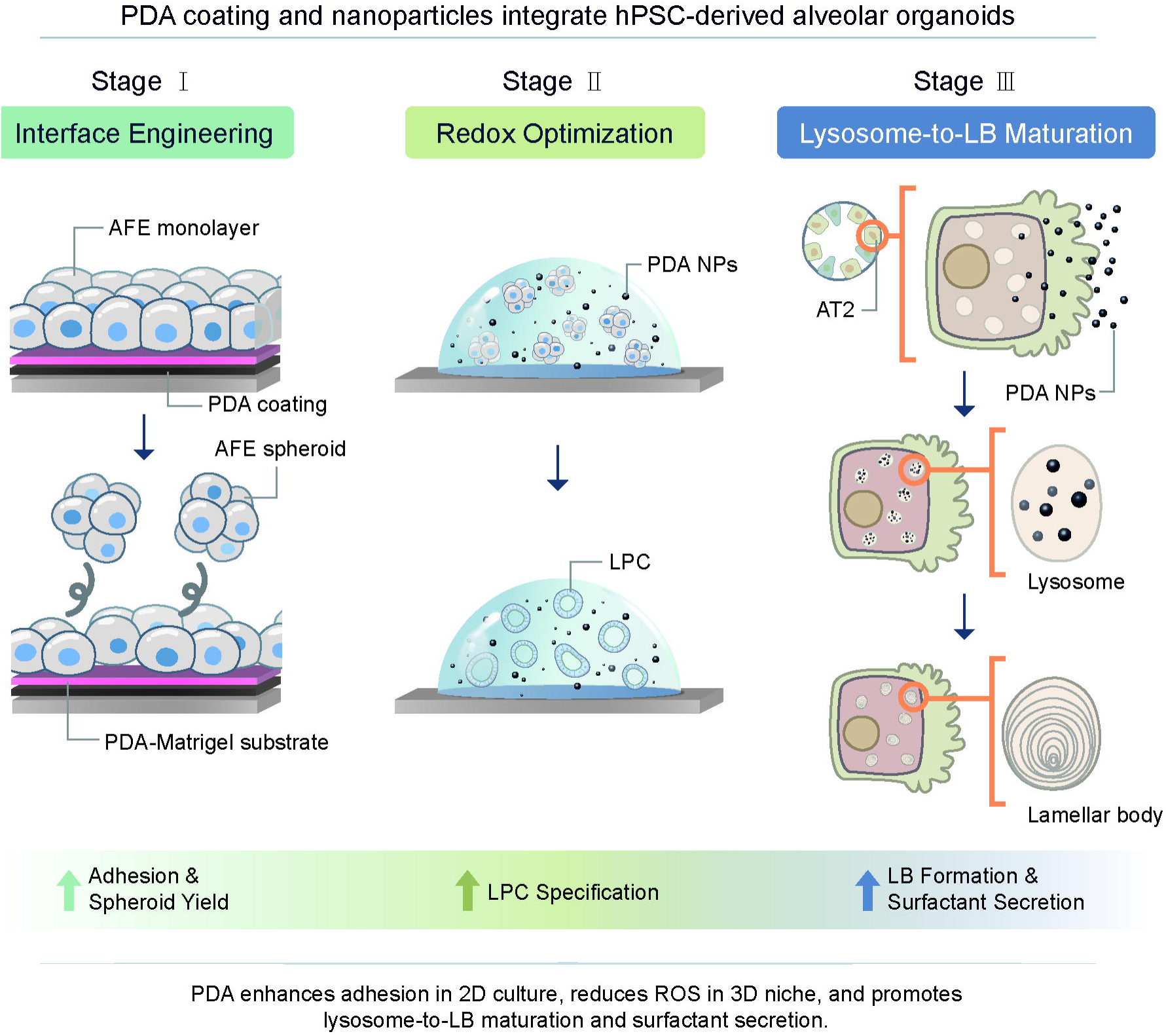

## 1. Introduction

Human pluripotent stem cell (hPSC)–derived alveolar organoids (ALOs) have emerged as powerful *in vitro* platforms for investigating human lung development[1], providing renewable sources of alveolar epithelial type 2 (AT2) for alveolar regeneration and repair[2], and accelerating respiratory drug discovery[3, 4]. Within the context of lung organoid engineering, hPSC-ALOs generation typically involves two-dimensional (2D), three-dimensional (3D) and suspension cultures[5–7]. The 2D phase facilitates efficient nutrient and oxygen exchange but the cell-culture substrate interactions should be carefully monitored to control differentiation outcomes. In contrast, 3D organoids more effectively recapitulate the *in vivo* epithelial cells reside microenvironment and cell-cell interactions[8]. However, the confined geometry and metabolic activity of organoids promote the accumulation of reactive oxygen species (ROS) both endogenous or exogenous[9–12]. Polydopamine (PDA), a mussel-inspired polyphenolic biomaterial, can be readily fabricated into substrate coatings and nanoparticles (NPs) with maintained antioxidant and bioadhesive properties[13, 14]. However, whether PDA could be incorporated into hPSC-ALOs system to enhance ALOs generation and the underlying mechanisms have not been investigated.

*In vitro* hPSC-ALOs differentiation follows key stages of *in vivo* embryonic lung development, progressing through definitive endoderm (DE), anterior foregut endoderm (AFE), lung progenitor cells (LPCs) and ALOs. There are many strategies from different labs to differentiate hPSC to ALOs[2, 6, 15, 16]. Our lab primarily cited the method from James Wells and Jason Spence and have published several papers[17–21]. During early differentiation, hPSCs are directed to form self-assembled AFE spheroids using a 2D monolayer culture format. Subsequently, self-assembled AFE spheroids were collected and embedded into Matrigel for ALOs differentiation. Efficient and consistent generation of AFE spheroids is critical for successful downstream organoid formation[16]. However, low AFE yield is common and may arise from suboptimal cell density, failed DE differentiation and insufficient activity of inductive growth factors[22]. Moreover, attempts to scale up AFE spheroids production often induced detachment of the monolayer, thereby limiting both the duration and overall efficiency of AFE spheroids harvest. Previous studies have reported that cell-mediated contractile forces could disrupt extracellular matrix (ECM) coatings on culture substrates, including culture plates and microfluidic surfaces[23]. While PDA coating significantly strengthens the interface between the ECM and underlying substrate, its impact on spheroids formation and detachment remains unexplored. In addition to enhancing cell adhesion, recent studies found PDA coating promotes cell migration by modulating integrin availability[24]. However, PDA coating itself led to less global migration yet more local motility of mesenchymal stem cells (MSCs)[25]. Therefore, whether incorporating PDA coating into the 2D culture system would enhance or dampen the production, stability, and yield of hPSC derived AFE spheroids warrants systematic investigation.

LPCs specification and early lung epithelial development rely on 3D ECM support. From previous publications, restricted nutrient diffusion, limited oxygen transport and metabolic waste accumulation within organoids and the surrounding 3D matrix often leads to elevated levels of ROS, which impair organoid growth, reduce differentiation efficiency and induce apoptosis[9, 26]. PDA NPs exhibit high colloidal stability and well-dispersibility in aqueous and biological media, enabling them to distribute uniformly within 3D hydrogels for organoid culture[27–29]. Indeed, PDA NPs have been widely incorporated into hydrogels to scavenge ROS in the microenvironment during cell regeneration[30–32]. Beyond their extracellular effects, PDA NPs can traverse the cell plasma membrane and are readily captured by lysosome and mitochondrial[33, 34]. Previous study reported supplementation of PDA NPs enhances mitochondrial metabolic activity in iPSC-derived cardiomyocytes by reducing intracellular ROS[35]. Lysosomes, in addition to their essential role in cellular degradation and homeostasis, contribute critically to the formation of lamellar body (LB), a hallmark feature of AT2 maturation and surfactant production[36, 37]. Although lysosomal accumulation of PDA NPs has been reported, their potential influence on lysosome-to-LB maturation dynamics and functional development during AT2 differentiation remains unexplored. Therefore, whether PDA NPs can support hPSC-ALOs differentiation and modulate key intracellular organelle processes to enhance AT2 maturation still need further investigation.

Here, we propose that PDA serves as a bioactive, stage-specific enhancer for hPSC-ALOs differentiation and maturation. Specifically, our results indicated PDA coatings during early adherent stages (hPSC, DE and AFE) strengthen ECM anchorage, reduce detachment losses, and improve the yield and duration of AFE. PDA NPs incorporated during 3D and suspension culture mitigate ROS accumulation, preserve epithelial viability, and potentiate maturation cues driving LPCs generation and AT2 functionality. This dual-phase (coating and NPs) design integrates PDA with intra- and extra-cellular redox regulation, addressing the key barriers of efficiency, robustness, and functional maturity. Through this strategy, we aim to engineer hPSC-ALOs with enhanced differentiation fidelity and physiological relevance by systematically using of PDA throughout the differentiation process.

## 2. Materials and methods

### 2.1 Materials

Dopamine hydrochloride (Aladdin, Cat#D103111), Tris (Sangon Biotech, Cat#A100826), Tris hydrochloride (Sangon Biotech, Cat#A100234), SYLGARD™ 184 Silicone Elastomer Kit (Dow, Cat#4019862), ammonia (Aladdin, Cat#A112081), 2,2-Diphenyl-1picrylhydrazyl (DPPH; TCI, Cat#D4313). The reagents and detailed medium formulations for each step of hPSC-derived ALOs differentiation are provided in the Table S1 and S2. Primer sequences and antibodies are provided in Table S3 and Table S4, respectively.

### 2.2 Fabrication of PDA-Matrigel coated substrates

Dopamine hydrochloride was dissolved in 10 mM Tris buffer (pH 8.5) at the concentration of 1 mg/mL and incubated with tissue culture plates for 2h at 37°C. Plates incubated with Tris buffer alone served as the control substrate. After coating, substrates were rinsed three times with deionized (DI) water and sterilized with UV light for at least 30 min. Matrigel was used to coat the surface for 2h at 37°C prior to cell seeding.

### 2.3 Substrate characterization

#### 2.3.1 Atomic force microscopy (AFM)

Substrate surface topography and mechanical properties were characterized using an AFM (Dimension FastScan, Bruker, USA). Non-conductive silicon nitride cantilevers (tip radius 10µm; spring constant 0.03 N/m; driving frequency of 10-20 kHz) were used for measurements with a typical scan rate of 1 Hz. Force volume (FV) mode was applied to characterize the modulus, surface morphology and roughness. Samples were fixed onto the AFM sample holder using an adhesive tape, and measurements were performed in a liquid environment at room temperature (RT). For each sample, three maps were acquired from different macroscopic regions. The root mean square roughness (Rq) was calculated using Version 1.8 Nanoscope analysis software (Bruker, USA).

#### 2.3.2 Water contact angle (WCA) measurement

WCA measurements were performed using a goniometer (SDC-200S, Sindin, China). PDA and PDA-Matrigel modified surfaces were dried under a cell culture hood for 30–60 min. WCA was measured using a 5μL DI water droplet at RT, and the angles were recorded after 5s. Three measurements per sample were averaged.

#### 2.3.3 Matrigel droplet retention assay

To assess the adhesive effect of PDA coating, an array of Matrigel droplets was prepared on the PDMS surfaces either treated by PDA or non-treated control. The substrates with Matrigel droplets were incubated in PBS solution and constantly agitated by a rotating shaker at 100rpm for 4h at RT. The number of remaining droplets was imaged and counted every 2h.

### 2.4 Synthesis of PDA NPs

PDA NPs were synthesized via the oxidative polymerization of the dopamine. Briefly, 80 mg of dopamine was dissolved in a mixture of water/ethanol (28 mL/12 mL) under stirring. Then 200 μL of ammonia (28wt%) was added to the water/ethanol mixture and stirred overnight. Formation of PDA NPs was indicated by the color change from light brown to black. The PDA NPs were collected by centrifugation at 12,000 rpm for 12 min (repeated 3-5 times), washed and then lyophilized.

### 2.5 Characterization of PDA NPs

#### 2.5.1 Scanning and Transmission electron microscopy (SEM/TEM)

PDA NPs were imaged by SEM (GeminiSEM 500, Carl Zeiss, Germany). PDA NPs were dispersed in DI water and drop-casted onto conductive carbon tape and sputter-coated with gold prior to imaging. For TEM (Talos L120C, Thermo Fisher Scientific, USA), PDA NPs were dispersed in DI water and deposited on a copper grid for imaging.

#### 2.5.2 Dynamic light scattering (DLS) and zeta potential analysis

The hydrodynamic diameter (Dh) and surface charge of PDA NPs suspension were meaured using a Zetasizer (Nano-ZS 90, Malvern Instruments, UK) with a detection angle of 90°. The zeta-potential measurements were conducted in DI water. All measurements were carried out at RT and repeated three times.

#### 2.5.3 Fourier transform infrared spectroscopy (FTIR)

Fourier transform infrared (FTIR) spectroscopy was used to characterize the chemical functional groups of dopamine and PDA NPs. Briefly, dopamine powder and lyophilized PDA NPs were submitted for FTIR measurement. For analysis, samples were gently pressed into a thin, uniform film. FTIR spectra were collected using an FTIR spectrometer (Nicolet^TM^ iS50 FTIR, Thermo Fisher Scientific, USA) in the midinfrared region of 4000–400 cm^-^¹, with a spectral resolution of 4 cm^-^¹ and 32 scans per sample. Background spectra were recorded prior to each run and automatically subtracted. The resulting spectra of PDA NPs were compared with those of free dopamine to assess changes in characteristic functional-group vibrations associated with polymerization.

#### 2.5.4 DPPH radical scavenging assay

The antioxidant activity of PDA NPs in water was evaluated by the DPPH assay. DPPH was dissolved in 95% ethanol (100 μM) and 200μL was added to each well of a 96-well plate. PDA NPs were added at various concentrations (0, 1.0625, 3.125, 6.25, 12.5, 25, 50, and 100 μg/mL) and incubated for 25 min in the dark. Absorbance was measured at 516 nm. Radical scavenging activity was calculated referring to a previous study[13].

### 2.6 hPSC maintenance

The H1 hPSC line was maintained in mTeSR1 medium (Stemcell, Cat#85850) on 6-well plates at 37 °C, 5% CO_2_, and 100% humidity. The medium was changed daily, and cells were passaged every 5–6 days at split ration of 1:40.

### 2.7 hPSC-ALOs differentiation and maturation

The hPSCs were differentiated into ALOs based on the previously published protocols[6, 16]. Briefly, hPSCs were dissociated into single cells using Accutase (StemCell, Cat#7920) and seeded at a density of 1.6×10^5^ cells/well in 24-well culture plates with Y-27 (10 µM; MCE, Cat#HY-10071). At 70∼80% confluence, hPSCs were treated with DE medium for 3 days, followed by AFE medium. During days of 2 to 7 of AFE induction, spheroids were formed and floated in the culture supernatant. Next, AFE spheroids were encapsulated into Matrigel (growth factor reduced, GFR; Corning, Cat#354230) to support 3D culture. Approximately 200 AFE spheroids were mixed with 25 µL Matrigel and placed at the center of each well of a 24-well plate. The 24-well plate was gently inverted to avoid the precipitation of spheroids and incubated at 37 °C for 15min to allow gelation. Solidified Matrigel droplets were then overlaid with LPCs medium, which was refreshed every other day for 7 days. Subsequently, medium was then switched to ALOs medium. Early ALOs were released from Matrigel using Cell Recovery Solution (Corning, Cat#354253) at day 7 and transferred to ultra-low attachment 24-well plates for suspension culture for an additional week.

### 2.8 Cell viability assay

Cell viability was assessed using the Cell Counting Kit-8 (CCK-8; MCE, Cat#HY-K0301) assay. Briefly, hPSCs were seeded on Matrigel and PDA-Matrigel substrates in 24-well plates at 1.6 × 10^5^ cells/well for 48 h. The absorbance at 450 nm was determined using a microplate reader (Multiskan Sky, Thermo Fisher Scientific, USA). The cell viability was calculated as (OD of cells on PDA /OD of cells on control) × 100 %.

### 2.9 qPCR analysis

The total RNA extraction (Qiagen, Cat#74106) of hPSC, DE, AFE, LPCs and ALOs, cDNA synthesis (Thermo Fisher Scientific, Cat#EP0753) and qPCR reactions (Takara, Cat#RR820A) were performed according to the manufacturers’ instructions. GAPDH was used as the housekeeping genes for normalization.

### 2.10 Immunofluorescence (IF) staining

IF staining was performed following published protocols[38]. Briefly, DE and AFE adherent cells were replated on Matrigel coated round cover slips and stained for confocal imaging. AFE spheroids, LPCs and ALOs were performed whole-mounting staining and then imaged on LSM-980 confocal microscope (Carl Zeiss, Germany).

### 2.11 Western Blotting (WB)

Total protein was extracted from LPCs and ALOs using RIPA lysis buffer (Beyotime, Cat#P0013B) supplemented with a protease inhibitor cocktail (Beyotime, Cat#P1006). Protein concentration was quantified using a BCA Protein Assay Kit (Beyotime, Cat#P0010). Equal amounts of total protein (30 μg) were loaded onto 4–20% precast polyacrylamide gels (GenScript, Cat#M42015C) for separation. Protein molecular weight markers (Invitrogen, Cat#26619) were added to determine the size of the protein bands. The intensity of WB bands was quantified using densitometry analysis. After acquiring the blot images, the grayscale values of the bands were measured using ImageJ software.

### 2.12 Intracellular ROS and superoxide measurement

Intracellular ROS and superoxide of LPCs and ALOs were measured using ROS/Superoxide Detection Assay Kit (Abcam, Cat#1027308-3). Briefly, LPCs and ALOs were incubated with oxidative stress detection reagent and superoxide detection reagent for 30min. Cell nucleus were stained with Hoechst 33342 (Thermo Fisher Scientific, Cat#62249). The organoids were then imaged on LSM-980 confocal microscope (Carl Zeiss, Germany).

### 2.13 Cellular uptake and lysosomal localization of PDA NPs

ALOs were incubated with FITC-labeled PDA NPs for 24 hours, followed by washing with ALOs medium. Then, lysosomes were labeled using 100 nM LysoTracker™ Red DND-99 (Thermo Fisher Scientific, Cat#L7528) for 30 minutesat RT in the dark. Nuclei and cell membranes were counterstained with Hoechst 33342 (Thermo Fisher Scientific, Cat#62249) and CellMask (Invitrogen, Cat#C10046) for 10 min. ALOs were then imaged using an LSM-980 confocal microscope (Carl Zeiss, Germany).

### 2.14 Live cell imaging of lamellar bodies in ALOs

ALOs were incubated in alveolarization medium with β-BODIPY FL C12-HPC (1 μM; Life Technologies; Cat#D-7392) for 24 h at 37 °C, 5% CO_2_. Organoids were washed and incubated with LysoTracker Red DND-99 (100 nM; Invitrogen, Cat#L7528) in alveolarization medium for 30 min at RT. After being washed twice, the samples were transfered to a confocal culture plate (15mm; Nest, Cat#801002) for imaging.

### 2.15 TEM of ALOs

TEM was used to analysis of intracellular morphological ultrastructures of organelles in ALOs associated with functional maturation. ALOs were collected and processed for TEM following previously described methods. Ultrathin sections were cut on Ultra-microtome (Leica EM UC7, Germany), mounted on grids, stained with uranyl acetate and lead citrate and imaged using a TEM equipped with a digital camera.

### 2.16 Bulk RNA sequencing (bulk RNA-seq) and bioinformatic analysis

AFE adherent cells and ALOs from Ctrl and PDA groups (3 biological replicates per group) were lysed with TRIzol™ reagent (Thermo Fisher Scientific, Cat#15596018) and flash frozen in liquid nitrogen for bulk RNA-seq. Quality testing, database construction and RNA-seq were subsequently performed by Annoroad Gene Technology (Beijing, China). The initial exploratory analysis, included PCA, volcano plots, heatmaps and bubble plots, were conducted using the SangerBox online platform (http://www.sangerbox.com/tool)[39]. Differential expression analysis identified significantly genes with P-value<0.05 and |log2(FC)| >1.5.

### 2.17 Statistical analysis

All the experiments were repeated at least three times. Data are presented as the mean ± Standard Deviation (SD). Statistical analyses were performed using one-way ANOVA with Tukey’s post hoc test or Student’s t test using GraphPad Prism 8.0.2 software. A P value < 0.05 was considered statistically significant.

## 3. Results

### 3.1 Stage-adaptive integration of PDA across hPSC-ALOs differentiation

In this study, the differentiation of hPSC into ALOs involved three distinct culture formats: 2D monolayer culture, 3D Matrigel-based culture, and suspension organoids culture (Fig. 1A). PDA was incorporated throughout the hPSC-to-ALOs differentiation process in multiple forms, including 2D surface coatings, nanoparticle–hydrogel composites, and nanoparticle-suspended culture media, enabling systematic functional investigations across all differentiation stages.

**Fig. 1.**
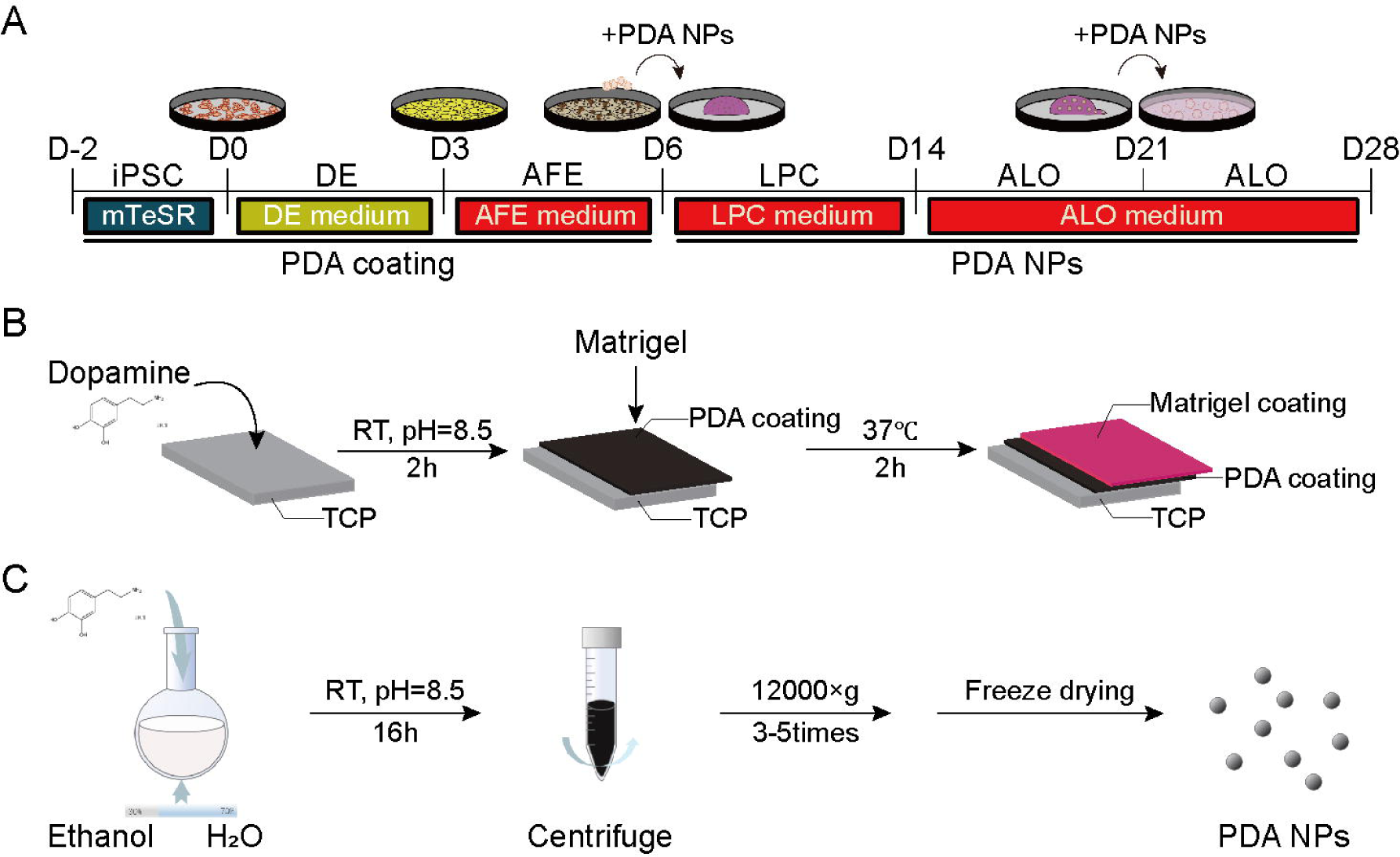
Experimental scheme of PDA coating and NPs application in hPSC-derived ALOs. A) Directed differentiation protocol from hPSC to ALOs with the use of PDA coating and NPs; B) Schematic illustration of the fabrication process of the PDA–Matrigel coating; C) Synthetic procedure of PDA NPs.

In the 2D monolayer stage, PDA was applied as a coating on tissue culture plates (TCP) to evaluate its impact on hPSC-AFE differentiation. PDA coating was achieved by incubating the plates surface with dopamine solution at pH 8.5 for 2 hours at 37°C, followed by the coating of Matrigel (Fig. 1B). The surface stiffness and roughness were analyzed by AFM on the substrates of TCP, PDA-coating TCP (PDA), Matrigel-coating TCP (Matri) and PDA-Matrigel-coating TCP (PDA-Matri) to examine the mechanical and morphological properties. Our results indicated the PDA coating significantly decreased the elastic modulus of TCP from 5.22±0.23 GPa (TCP) to 4.69±0.05 GPa (PDA) (P<0.01) and Matrigel from 136.30±10.66 kPa (Matri) to 38.00±7.48 kPa (PDA-Matri), respectively (Fig. S1A, B). Furthermore, the PDA coating significantly increased surface roughness compared to TCP and Matrigel substrates (Fig. S1C, D). Taken together, PDA coating significantly increased the roughness but decreased the elastic modulus which suggested a more cell attachment and culture favorable microenvironment.

WCA measurements were performed to evaluate the surface wettability of the substrates. Compared to TCP (88.3±1.2°), PDA-coated substrates exhibited a significantly decreased WCA (59.1±1.3°), indicating enhanced surface hydrophilicity. PDA-Matri further decreased the WCA from 41.7±5.2° (Matrigel-coated surface) to 24.4±0.9°, suggesting the PDA coating enhanced Matrigel spreading and hydrophilicity (Fig. S1E). To determine the bioadhesive property of the PDA coating, an array of hydrogel droplets was added in the culture plates and continuously agitated at 100rpm for 4h based on previous study[23](Fig. S1F). Under this turbulent condition, the hydrogel droplets array on the PDA substrate adhere more tightly without detachment for at least 2 hours. In contrast, 89% of the Matrigel droplets detached from the control substrate within 2h (Fig. S1G, H), suggesting PDA coating promoted hydrogel adhesion. In summary, the PDA coating was made as an intermediate layer between TCP and Matrigel coating to enhance Matrigel adhesion and spreading properties.

During the organoids (3D) differentiation stages, PDA was fabricated into NPs and mixed either with the Matrigel matrix or suspension medium. PDA NPs were prepared by dopamine self-polymerization following previous publications[40]. Briefly, dopamine was dissolved in a mixture of water/ethanol at pH8.5 and stirred overnight at room temperature. The solution was then centrifuged at 12,000 g and freeze-dried to obtain the PDA NPs (Fig. 1C). FTIR analysis confirmed the successful synthesis of PDA NPs, as evidenced by a characteristic quinone peak at 1700 cm^-^¹ and a broadened O–H stretching band at 3200–3400 cm^-^¹ due to extensive hydrogen bonding, which are distinct features of PDA compared with DA (Fig. S2A). The size and morphology of the PDA NPs were analyzed using SEM and TEM. The particle size of PDA NPs is 111.7±12 nm (Fig. S2B, C). DLS results indicate that the PDA NPs can be well dispersed in water with the size distribution at 198.3±18.5 nm (Fig. S2D). The zeta potential of PDA NPs is around -30 mV (Fig. S2E). DPPH was used to assess the antioxidant activities of the PDA NPs. The results showed that PDA NPs can effectively scavenge ROS, with the antioxidant efficiency increased with the increasing concentration (Fig. S2F). Taken together, PDA was fabricated in different formats (coatings and NPs) and stage-adapted throughout the hPSC-to-ALOs differentiation process, with enhanced interface adhesive and retained antioxidant properties.

### 3.2 PDA coating enables scalable and prolonged AFE spheroids production

The 2D culture covers the differentiation stages from hPSC to AFE. PDA coating has been widely applied in 2D culture plates modification. However, the PDA coating in hPSC differentiation into AFE would be a beneficial or hinder in AFE spheroids formation has not been studied.

We firstly examined the effect of PDA coating on hPSC stemness maintenance. The bright-field images of hPSC revealed the rounded hPSC colony adhere and proliferate normally without morphological differences between control and PDA treated substrate after 48 hours (Fig. S3A). CCK-8 assay was performed to assess the effect of PDA on hPSC viability, which indicated PDA coating did not affect cell viability (Fig. S3B). The stemness gene expressions of OCT4 were similar between the hPSC cultured on control and PDA coating substrate (Fig. S3C). Previous study has reported PDA coating enhanced MSCs stemness through eliminating the ROS accumulation in the MSCs[41]. We did not find the enhanced stemness in our hPSC is likely due to the stem cells only maintained 48 hours and then entered DE differentiation.

DE differentiation lasted for 3 days. Bright-field images of DE cells revealed similar morphological characteristics on Matrigel and PDA-Matrigel substrate (Fig. S3D). The expression levels of SOX17 and FOXA2 showed no significant differences between cells cultured on Matrigel and PDA-Matrigel substrate (Fig. S3E). IF staining of SOX17 and FOXA2 revealed that DE cultured on PDA coating exhibited similar SOX17 positive (green) and FOXA2 positive (red) cells compared with the control (Fig. S3F, G). These results indicated the PDA coating did not interfere the induction of DE cells. Taken together, PDA coating maintained the hPSC and DE cells viability and differentiation properties.

During AFE stage, self-assembled spheroids formed, detached from cell monolayer, and floated in culture medium, which were collected for further differentiation. During the AFE spheroids formation and de-attachment, we found cells cultured on Matrigel exhibited about 60% cell monolayer pealed from the culture plate (Fig. 2A, B). Previous study has reported PDA coating could enhance the interaction between the hydrogel coating and the culture plate[23]. Consistently, our result also found the PDA treatment significantly increase the adhesion of cell monolayer with the substrate and decrease the ratio of detachment to 15% (Fig. 2B). The AFE monolayer gene expression of FOXA2 was not affected by PDA coating (Fig. 2C). IF staining of FOXA2 in AFE monolayer showed PDA maintained AFE differentiation efficiency (Fig. 2D). AFE spheroids generation typically initiates around day 2, peaks during days 3-4, and then rapidly declines by day 5 as AFE monolayer begins to peel, leading to failure of further spheroid production (Fig 2E). However, PDA coating significantly increased spheroids production and extended the spheroids generation period from day 2 to day 7 (Fig. 2E, F). The total number of spheroids increased from 220 /well to 360 /well of 24-well culture plate (Fig. 2G). The gene expressions of FOXA2 in AFE spheroids were not affected by PDA coating (Fig. 2H). Whole-mounting IF staining confirmed spheroids obtained from PDA-Matrigel substrate displayed similar homogeneous expression of FOXA2 (red) (Fig. 2I). Together, these findings demonstrate that PDA coating reinforces AFE monolayer cell-substrate adhesion to prevent the monolayer peeling induced failure of AFE spheroids collection, yields more AFE spheroids for extended period, without the change of the AFE cell fates.

**Fig. 2.**
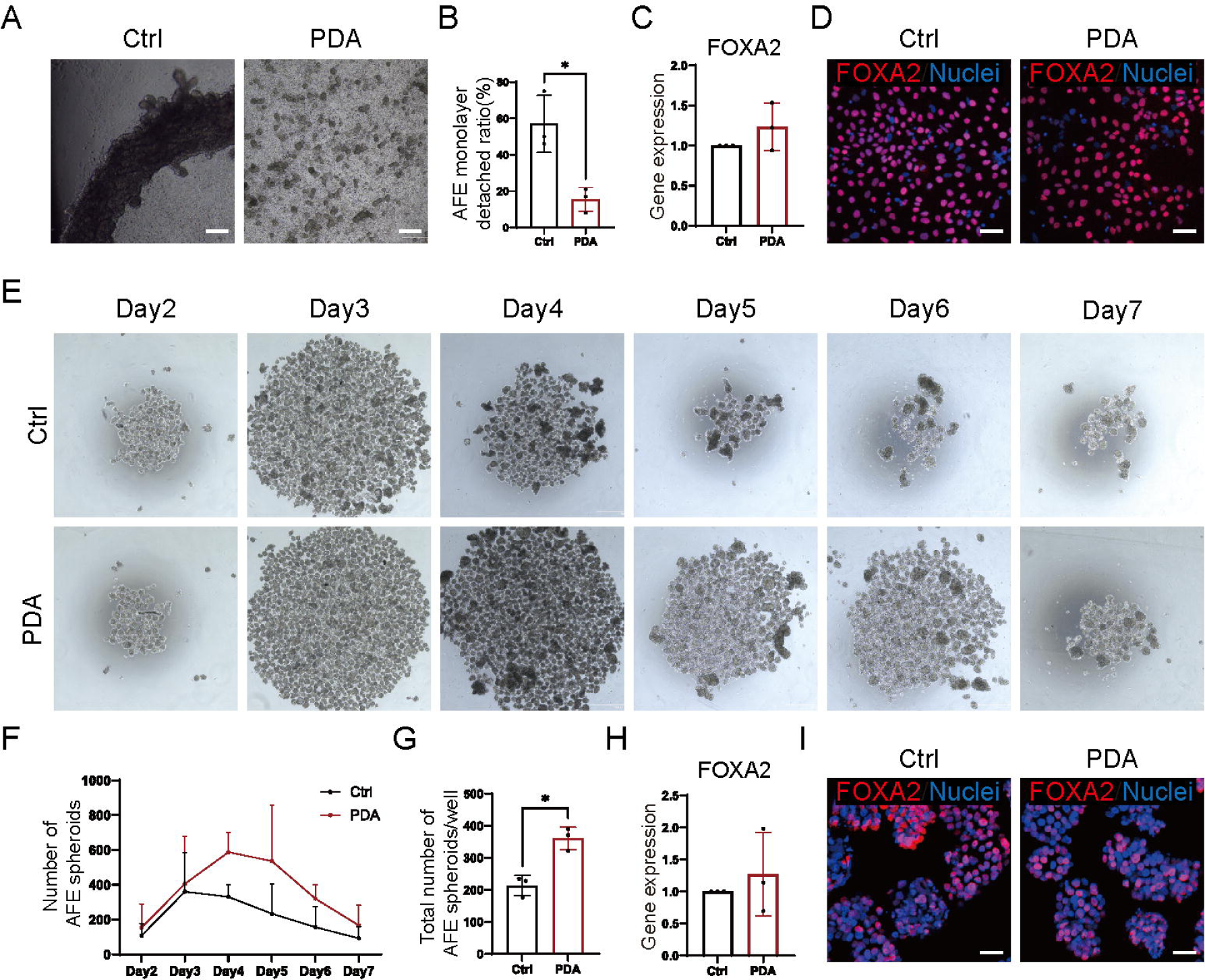
PDA coating reduces AFE monolayer detachment and boosts AFE spheroids yield and duration. A) Bright-field images of AFE adherent cells showing monolayer detachment from control (untreated culture plate, ctrl) but non-detachment from PDA-treated culture plate (PDA) (scale bar: 100µm); B) Quantification of AFE monolayer detachment ratio in different coating conditions; C) qPCR analysis of FOXA2 expression of AFE adherent cells; D) IF staining of FOXA2 of AFE monolayer cells (scale bar: 50µm); E) Representative bright-field images of AFE spheroids collected from day 2 to day 7 of AFE stage; F) Quantitative analysis of the number of AFE spheroids from day 2 to day 7; G) Quantitative analysis of the total number of AFE spheroids; H) qPCR analysis of FOXA2 expression of AFE spheroids under control and PDA treatment; I) Immunofluorescence staining of FOXA2 in AFE spheroids under control and PDA treatment (scale bar: 50µm).

### 3.3 PDA coating primarily modulates cell-cell and cell-ECM interactions at AFE

Bulk RNA-seq analysis was performed to elucidate the signaling pathways and molecular mechanisms underlying the effects of PDA-coated culture surfaces on AFE spheroids formation. The principal component analysis (PCA) demonstrated robust separation of the two groups, indicating that PDA coating induces a reproducible global shift in the AFE transcriptome (Fig. 3A). Among these, 215 significantly differentially expressed genes (DEGs; 98 upregulated and 117 downregulated) were identified in PDA-Matrigel group. The differential expression analysis identified substantial transcriptional reprogramming in response to PDA, with both up- and down-regulated gene sets (Fig. 3B). Unsupervised clustering of representative DEGs revealed a clear segregation between PDA and control samples, with high intra-group concordance (Fig. 3C). Notably, PDA exposure was associated with marked alterations in morphogenesis, cell adhesion, ECM organization, and developmental signaling, including genes related to extracellular and matrix-related components compartments (including COL4A4, COL11A2, PRSS1, INHBE, BMP5 and WNT5A), together with multiple transcriptional regulators and signaling mediators implicated in AFE patterning and epithelial development (including PIK3R5, EGF, TGFB2, MAPK8IP1, CYP26C1, FGF17, and FGF20).

**Fig. 3.**
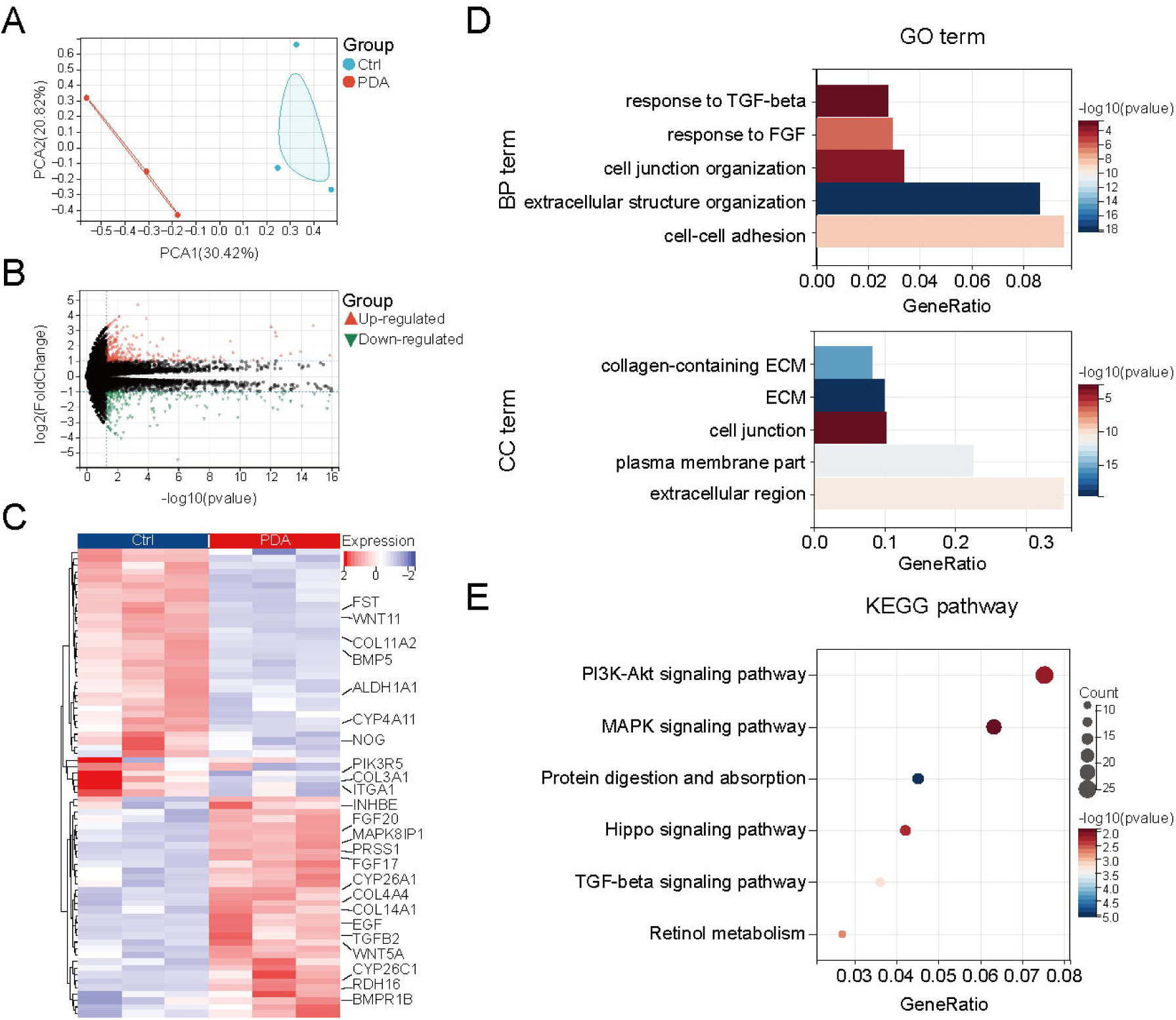
Bulk-RNA seq results indicate PDA coating primarily modulates cell-cell or cell-ECM interactions at AFE stage. A) PCA showing separation between Ctrl and PDA; B) Volcano plot of DEGs between Ctrl and PDA; C) Heatmap illustrating the upregulation and downregulation of Ctrl to PDA; D) Selection of significantly enriched GO terms in BP and CC categories; E) Selection of significantly enriched KEGG pathways.

The AFE monolayers on the PDA coating were enriched for genes with GO biological process (BP) terms related to responses to FGF and TGF-β signaling, extracellular structure organization, cell–cell adhesion, and cell junction organization (Fig. 3D). Consistently, cellular component (CC) terms were enriched for ECM, collagen-containing ECM, extracellular region, cell junction, and cell–ECM interfaces by PDA treatment. KEGG analysis further identified enrichment in PI3K–Akt, MAPK, TGF-β, Hippo, retinol metabolism and protein digestion and absorption signaling pathways (Fig. 3E). Among these signaling pathways, PI3K–Akt, MAPK are down-stream cascades of FGFR, while TGF-β and Hippo signaling pathways integrate growth factor and mechanotransduction cues to promote epithelial morphogenesis and spheroids formation. Collectively, these transcriptomic signatures suggest that PDA promotes AFE morphogenesis by strengthening cell-cell, extracellular programs and engaging growth factor-linked signaling, consistent with enhanced spheroid generation without altering AFE identity.

### 3.4 PDA NPs eliminate ROS accumulation in 3D Matrigel and promote NKX2-1^+^ LPCs generation

AFE spheroids were encapsulated in Matrigel for subsequent differentiation into LPCs. Previous study has reported that 3D hydrogel droplets can accumulate ROS within the microenvironment, thereby inducing organoids apoptosis[9]. To mitigate this effect, we incorporated PDA NPs with antioxidant property into the Matrigel (Fig. 4A). The bright field images showed single cells developed into LPCs organoids by day 7 (Fig. 4B). Importantly, the incorporation of PDA NPs into Matrigel did not affect LPCs morphogenesis. Live-cell staining revealed substantially reduced levels of ROS and superoxide in PDA-treated LPCs, which was further confirmed by quantitative fluorescence analysis (Fig. 4C, D). The PDA NPs were anchored in Matrigel matrix without entering the LPCs organoids. Therefore, the significant reduction of ROS in the LPCs is likely by effectively exert antioxidant property in the ECM microenvironment, thereby eliminating ROS within LPCs.

**Fig. 4.**
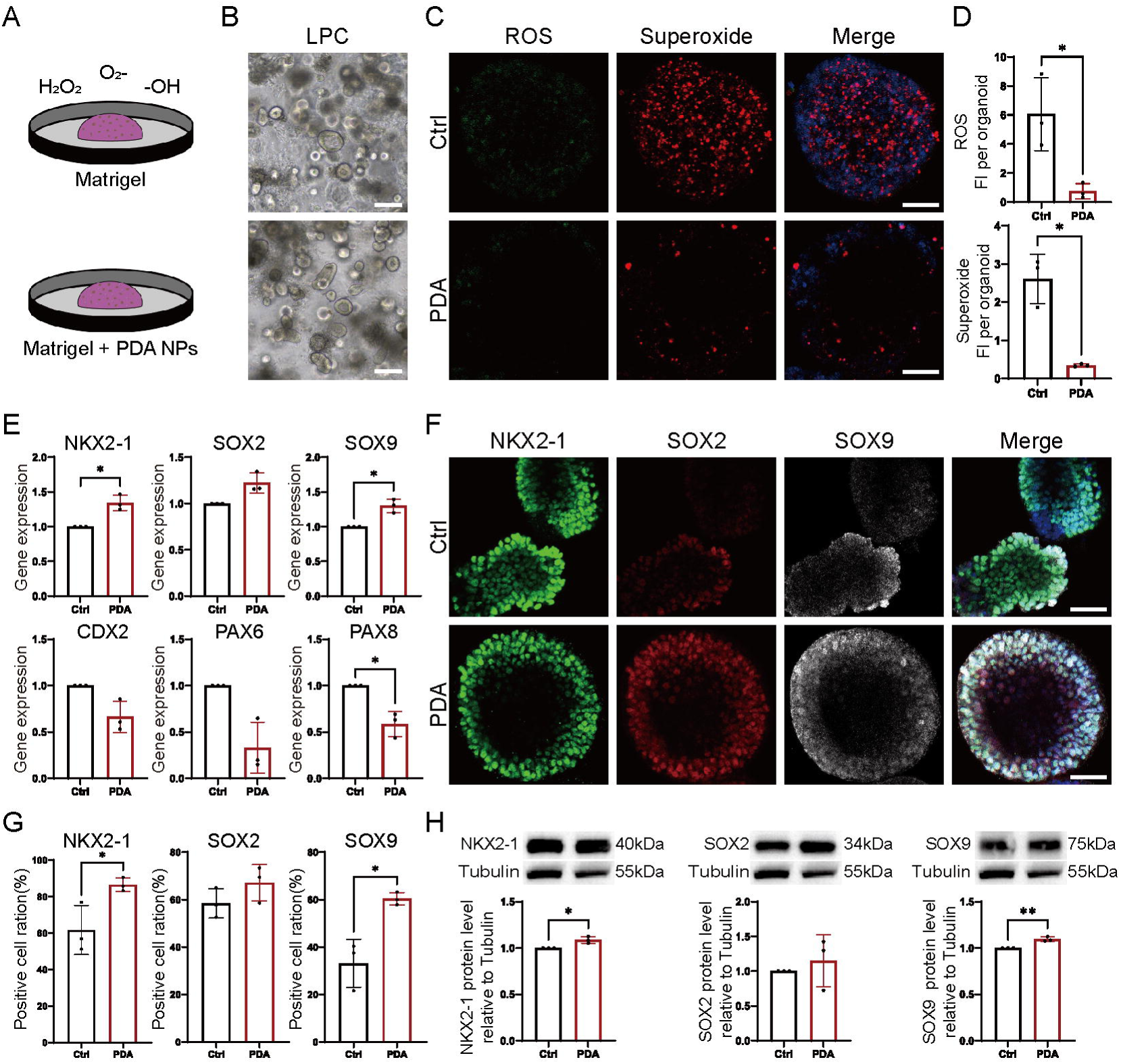
PDA NPs composed Matrigel promotes NKX2-1^+^ LPCs generation. A) Schematic representation of LPCs 3D culture: Matrigel and PDA NPs composite Matrigel; B) Bright-field images of LPCs. (scale bar: 200µm); C, D) Live-cell staining and quantification of intracellular ROS and superoxide levels in LPCs. (scale bar: 50µm); E) qPCR analysis of lung specific lineage-(NKX2-1, SOX2, and SOX9) and non-lung lineage markers (CDX2, PAX6, and PAX8) in LPCs; F, G) IF staining and quantitative analysis of NKX2-1, SOX2, and SOX9 expression in LPCs (scale bar: 50µm); H) Western blot analysis of NKX2-1, SOX2 and SOX9 protein levels.

The impact of PDA NPs on LPCs specification was then examined. qPCR analysis revealed that PDA treatment significantly upregulated LPCs markers NKX2-1 and SOX9, and reduced other lineage markers especially PAX8 (thyroid) with significance (P<0.05) (Fig. 4E). Whole-mounting IF indicated the NKX2-1(green), SOX2 (red) and SOX9 (white) were highly expressed in PDA-treated LPCs (Fig. 4F). The positive cell ratio of NKX2-1 and SOX9 were significantly increased in PDA-treated LPCs relative to the control (Fig. 4G). The proportion of NKX2-1 positive cells increased from 47% to 66%. Consistently, WB analysis demonstrated higher NKX2-1 and SOX9 protein levels in PDA-treated cells (Fig. 4H), indicating that PDA NPs effectively promote LPCs lineage specification.

These findings indicated that PDA NPs doped in Matrigel droplets mitigate oxidative stress and enhance NKX2-1^+^ LPCs generation in this 3D culture environment, suggesting PDA provides a more favorable redox environment for LPCs development.

### 3.5 Suspended PDA NPs reduce ROS and promote AT2 maturation

NPs could be internalized via endocytosis, captured by intracellular organelles, and involve in cell differentiation and function[35, 42]. We therefore investigated whether PDA NPs could be endocytosed and subsequently influence alveolar differentiation and maturation. To this end, the early-stage ALOs (week 1) were released from Matrigel and cultured in suspension with PDA NPs to assess their effects on ALOs differentiation (Fig. 5A). The PDA NPs supplemented culture medium was refreshed every other day, and PDA NPs were observed to localize around and on the surface of ALOs (Fig. 5B). Oxidative stress was assessed to evaluate the intracellular redox-regulating capacity of PDA. Live-cell staining revealed markedly reduced levels of ROS and superoxide in PDA-treated ALOs (Fig. 5C). The quantitative fluorescence analysis showed a significant decrease in overall ROS/superoxide intensity per organoid (Fig. 5D).

**Fig. 5.**
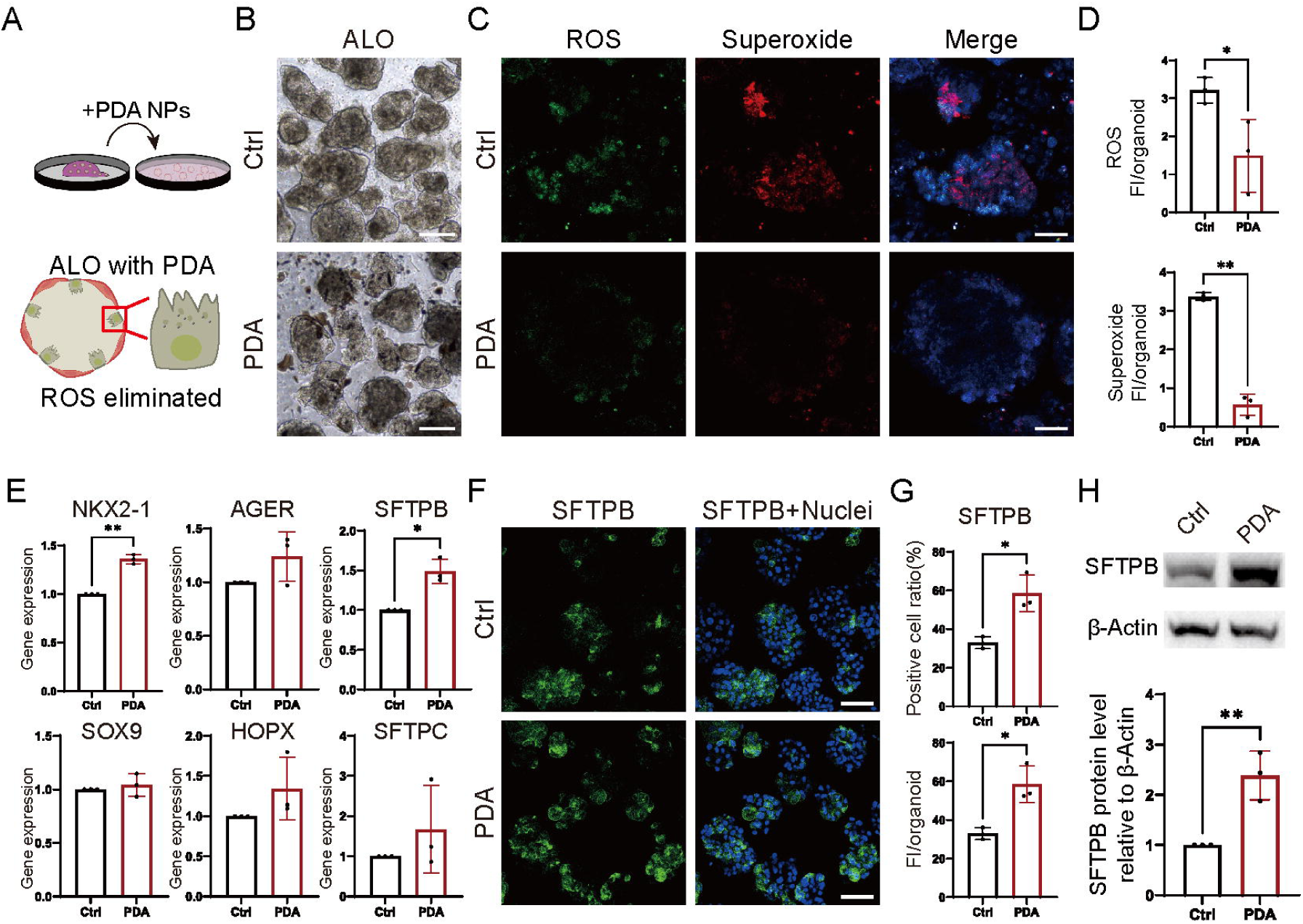
PDA NPs enhances AT2 maturation in lung organoids. A) Schematic representation of ALOs suspension culture; B) Bright-field images of ALOs. (scale bar: 200µm); C, D) Live-cell staining and quantification of intracellular ROS and superoxide levels in ALOs (scale bar: 50µm); E) qPCR analysis of ALOs marker genes (lung lineage: NKX2-1, SOX9; AT1: AGER, HOPX; AT2: SFTPB, SFTPC); F, G) IF staining and quantitative analysis of SFTPB expression in ALOs (scale bar: 50µm); H) Western blot analysis of SFTPB protein level.

The qPCR analysis demonstrated that PDA treatment significantly increased SFTPB expression whilst the expression of SFTPC, NKX2-1, SOX9, AGER and HOPX indicated elevated trend without statistical significance (Fig. 5E). IF staining further verified a significant increase in the proportion of SFTPB positive cells in PDA-treated ALOs, rising from 32% to 58% compared with the control (Fig. 5F, G). Consistently, WB analysis revealed a pronounced increase in SFTPB protein levels in the PDA-treated group compared to the control (Fig. 5H).

Together, these results indicate that during ALOs differentiation, PDA NPs significantly increase SFTPB expression and alleviate oxidative stress. Previous genetic and mechanistic studies have established that SFTPB is essential for LB biogenesis and maturation[43]. As LB development and maturation are hallmarks of AT2 functional maturation[44]. AT2 is the major alveolar epithelial stem cell[45]. We next investigated whether PDA NPs directly participate in LB biogenesis and maturation.

### 3.6 PDA NPs drive lysosome-to-LB maturation and surfactant secretion

LBs are specialized lysosome-related organelles in AT2 cells, whose biogenesis and maturation are tightly linked to the endolysosomal pathway. PDA NPs were recognized as invasive particles which were captured and decomposed by lysosome. We were curious whether PDA NPs were captured by lysosome in premature AT2 and participated in lysosome-to-LB maturation and function.

To investigate if PDA NPs could be captured by lysosome, FITC-labeled PDA NPs were added 24 hours before imaging. The fluorescent imaging indicated the co-localization of FITC-labeled PDA NPs (green) and lysosomes (red) within the ALOs (Fig. 6A). Approximately 70% of cells in ALOs contained detectable lysosomes, and among these lysosome-positive cells, about 60% of lysosomes exhibited co-localization with PDA NPs, suggesting efficient lysosomal uptake of PDA NPs within the organoids (Fig. 6B). Then, we assessed the secretion of pulmonary surfactant from LBs. PDA-treated ALOs exhibited increased β-BODIPY-positive lipid droplets and lysotracker positive lysosomes (Fig. 6C). What’s more, the co-localization of β-BODIPY and lysotracker also increased with the PDA treatment. The co-localization of β-BODIPY-positive lipid droplets and lysotracker indicates enhanced lysosome–lipid droplet interactions, implies the maturation of LBs[46]. This suggests enhanced formation of surfactant-associated storage organelles, a characteristic feature of mature AT2 cells. TEM further verified more lipid droplets were found in PDA-treated ALOs (Fig. 6D, yellow arrows), alongside the presence of internalized PDA NPs (Fig. 6D, blue arrows). ATP-binding cassette transporter (ABCA3), a marker for LBs, was co-stained with SFTPB to investigate the effect of PDA on AT2 maturation. IF stain indicated the number of ABCA3 positive cells and the co-positive cell ratio of SFTPB and ABCA3 increased in PDA-treated ALOs (Fig. 6E, F). TEM indicated PDA-treated ALOs displayed a higher density of mature LBs, identified by their characteristic concentric whorled membranes (Fig. 6G, red arrows). Collectively, these results demonstrate that PDA promotes AT2 cell maturation by enhancing the formation of LBs and the synthesis of essential surfactant components through PDA involvement in lysosome-to-LBs developmental process.

**Fig. 6.**
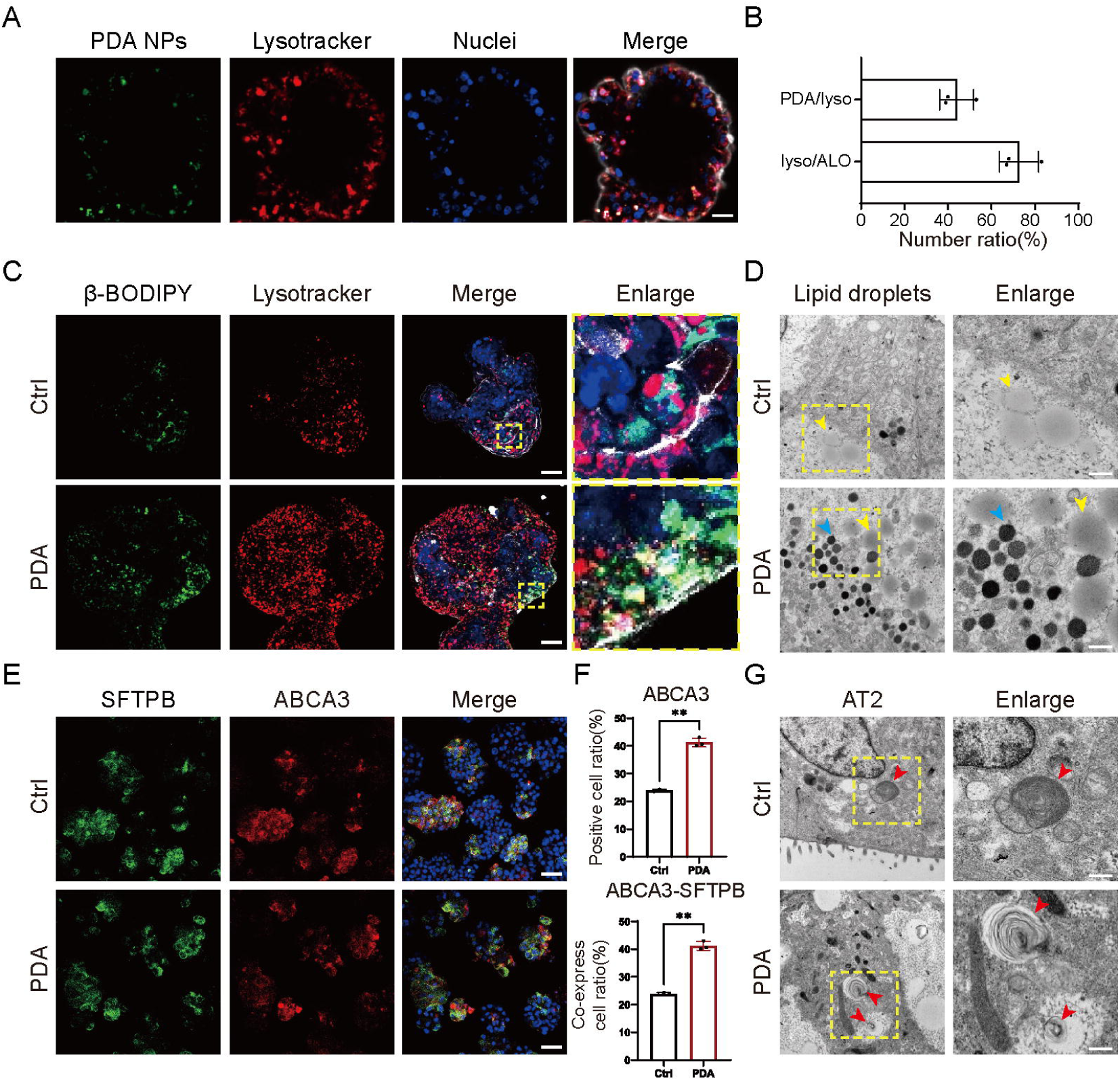
PDA NPs are internalized into ALOs, involves LB maturation and promotes surfactant secretion. A) Co-IF stain of FITC-PDA NPs and lysotracker in ALOs (scale bar: 20µm); B) Quantitative analysis of lysosomal uptake of PDA NPs in ALOs; C) Staining of ALOs using β-BODIPY to visualize AT2-secreted surfactant-associated lipid and Lysotracker to label lysosomes (scale bar: 50µm); D) TEM images of ALOs showing lipid droplets (yellow arrow heads), and PDA NPs (blue arrow heads) (scale bar: 100nm); E, F) IF staining of SFTPB and ABCA3, with quantification of ABCA3^+^ and ABCA3^+^SFTPB^+^ cell populations (scale bar: 50µm); G) TEM images of ALOs showing lamellar bodies (red arrow heads) (scale bar: 100nm).

### 3.7 PDA NPs activate endocytic-phagosome and epithelium maturation pathways in ALOs

To assess the global transcriptomic changes in gene expression and its underlying mechanism in PDA-treated ALOs, we performed bulk RNA-seq at the maturation stage in PDA-treated and control ALOs. PCA confirmed distinct clustering of the two groups with good within-group consistency (Fig. 7A). A total of 784 significantly DEGs (246 upregulated and 538 downregulated) were identified in PDA-treated ALOs. The distribution and up/downregulation of these DEGs are further illustrated in a volcano plot (Fig. 7B). Unsupervised clustering of representative differentially expressed genes revealed a clear segregation between PDA and control samples, indicating a robust transcriptomic shift induced by PDA NPs (Fig. 7C). PDA exposure regulated gene modules associated with epithelial secretory programs, xenobiotic/retinoid metabolism and redox-related enzymes (including WNT11, COL6A3, NKD1, CYP1A1, CYP1B1 and GHRL) (Fig. 7C).

**Fig. 7.**
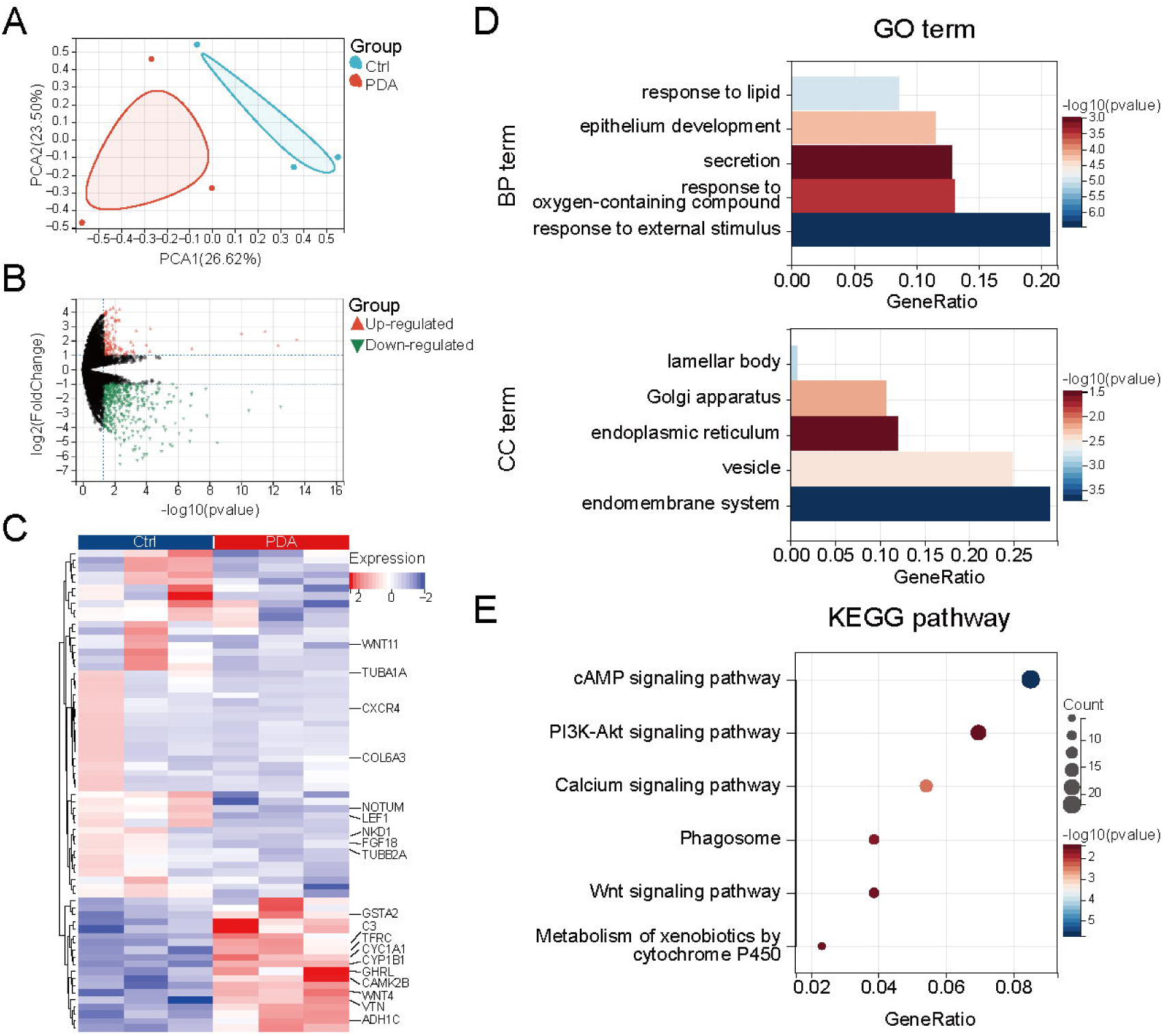
Bulk-RNA seq results demonstrate PDA NPs promote secretion and endocytosis-related pathways in ALOs. A) PCA showing separation between Ctrl and PDA. B) Volcano plot of DEGs. C) Heatmap illustrating the proximity of Ctrl to PDA. D) Selection of significantly enriched GO terms in BP and CC terms. E) Selection of significantly enriched KEGG pathways.

GO BP enrichment in PDA-NP-treated ALOs highlights responses to external stimulus was related to the internalization of PDA NPs (Fig. 7D). Enrichment of response to lipid, and secretion suggests AT2 maturation-related lipid secretory. Response to oxygen-containing compound aligns with the antioxidant property of PDA. Epithelium development supports the maturation of AT2. GO CC terms were dominated by the endomembrane system (vesicles, endoplasmic reticulum, Golgi apparatus) with additional enrichment of LB, indicating strengthened vesicular trafficking and LB organelle maturation. KEGG analysis further demonstrated that PDA-regulated genes were enriched in phagosome and metabolism of xenobiotics by cytochrome P450 indicated PDA NPs were endocytosed and metabolized (Fig. 7E). In addition, enrichment of cAMP, PI3K-Akt, calcium and Wnt signaling pathways, consistent with AT2 epithelium maturation. Taken together, bulk-RNA seq indicates that PDA NPs drive a coordinated maturation of AT2 and lysosome-to-LBs through reinforced endocytic-phagosome and epithelium development pathways.

In summary, PDA enhances hPSC-ALOs differentiation through a stage-specific, multi-level mechanism. Stage I (2D interface engineering, from hPSC to DE to AFE): PDA surface coating increases hydrophilicity and roughness and improves Matrigel spreading/adhesion, stabilizing the AFE monolayer, reducing monolayer peeling, and enabling scalable, prolonged production of AFE spheroids without altering AFE fate markers. Stage II (3D redox buffering, from AFE to LPCs): PDA NPs incorporated into Matrigel reduce ROS/superoxide accumulation, supporting NKX2-1^+^/SOX9^+^ lung progenitor specification and efficient progression toward alveolar fate. Stage III (maturing ALOs): PDA NPs are endocytosed and accumulate in lysosomes, enhancing endolysosomal trafficking linked to LB biogenesis, increasing ABCA3^+^/SFTPB^+^ LB maturation and promoting surfactant storage/secretion. Bulk RNA-seq further supports coordinated activation of secretory and endomembrane/endocytic programs during maturation.

## 4. Discussion

Although differentiation platforms for hPSC-ALOs are increasingly established, most engineering strategies primarily focus on improving reproducibility and reducing inter-organoid heterogeneity, while cell fate regulation remains largely dependent on developmental signaling cues and spatiotemporal administration of growth factors. Here, we introduce a material-enabled differentiation strategy that exploits the multi-staged processability and intrinsic biofunctionality of the mussel-inspired biomaterial PDA. By integrating PDA throughout the entire differentiation trajectory in the form of surface coating and NPs, we demonstrate stage-specific and complementary regulatory effects. PDA coating enhances cell–substrate adhesion and increase AFE yield during early adherent differentiation. In 3D Matrigel cultures, PDA NPs optimize the local microenvironment by reducing oxidative stress and promoting LPCs generation. During suspension-based alveolar maturation, PDA NPs are internalized by organoids and facilitate lysosome–LB biogenesis, thereby enhancing AT2 differentiation and functional maturation. Together, this work establishes a versatile material-assisted hPSC-ALOs platform that extends fate control beyond soluble growth factors-based regulation.

Efficient formation of AFE spheroids is a critical morphogenetic step in ALOs differentiation, yet its regulation remains incompletely understood. Previous studies have primarily attributed successful foregut spheroid initiation to biochemical patterning cues, particularly the coordinated activation of WNT and FGF signaling pathways[47]. In contrast to these chemically driven strategies, we identify PDA coating as a material-based regulator of AFE spheroid initiation that rather than directly engaging WNT or FGF signaling, PDA primarily modules mechanosensitive pathways such as PI3K–Akt, MAPK, Hippo, and TGF-β, which are widely recognized as signals that integrate extracellular adhesion cues with cytoskeletal dynamics and tissue integrity[48]. PDA also stabilizes adherent AFE monolayers through ECM deposition and remodeling protein digestion and absorption, likely represents adaptive responses to changes in cell–substrate coupling that facilitate collective detachment and spheroid cohesion. By applying PDA coating during the spheroid generation phase of adherent cell differentiation, this strategy leverages material-driven regulation of cell-substrate interactions to improve spheroids self-organization. Therefore, PDA promoted AFE spheroids formation through primarily mechanical pathways, ECM deposition and metabolism without modulating fate-specific pathways.

PDA coatings and NPs possess well-established antioxidant properties; however, the persistence of these effects during multi-stage organoid differentiation remains unclear. Previous studies generally evaluated PDA-mediated redox regulation within a short time (typically 24–72 h), during which significant reductions in intracellular ROS levels were consistently observed in stem cell cultures and nanoparticle-based systems[35, 41]. In our study, redox-related pathways were not enriched during early differentiation stages (AFE) cultured on PDA-coated substrates (Fig. 3D). This likely reflects the catechol and semi-quinone groups on PDA coatings can be gradually oxidized or consumed during early cultivation with daily medium exchange[41]. Moreover, AFE induction represents the late stage of endoderm differentiation which required moderate oxidative stress for early developmental patterning rather than limiting[49]. Thus, PDA coating at the AFE stage may primarily retain its bioadhesive and mechanical functions, while its antioxidant capacity is attenuated and no longer rate-limiting for cell behavior. In contrast, PDA NPs show pronounced antioxidant effects in LPCs (Fig. 4C) and ALOs stages (Fig. 5C), when metabolic demand and diffusion constraints substantially increase oxidative burden[50]. Unlike surface-bound coatings, PDA NPs function within the ECM and following cellular uptake, enabling sustained scavenging of ROS in both extracellular and intracellular compartments[35, 51]. The cell culture medium exchanged every other day which also elevated ROS accumulation and augmented antioxidant properties of PDA NPs. Consistently, we observed elimination of ROS and enrichment of oxidoreductase activity related genes by PDA NPs. Functionally, PDA NPs mediated ROS reduction to generate a more favorable microenvironment for progenitors and alveolar maturation.

In our study, we observed that PDA NPs were internalized by ALOs and predominantly accumulated within lysosomes, consistent with prior reports describing lysosomal sequestration of PDA-based nanomaterials[34, 35]. Furthermore, PDA NPs promoted the functional maturation of AT2 cells in ALOs, as evidenced by enhanced LB formation and increased surfactant secretion (Fig. 6). Previous studies have demonstrated that retinoid signaling and lysosomal lipid metabolism are integral to AT2 differentiation and surfactant homeostasis, and that perturbations in these processes impair LB formation and alveolar function[52, 53]. In this context, the observed activation of retinol metabolism, oxidoreductase activity, and secretory vesicle–associated programs in PDA-treated ALOs are consistent with enhanced lysosome-to-LB functional maturation. Such lysosome-centered adaptive responses to persistent metabolic or xenobiotic stimuli have been shown to drive organelle functional reprogramming, supporting the plausibility of a compensatory maturation mechanism that enhances LB biogenesis and surfactant synthesis and secretion in AT2 cells[54, 55]. Based on these findings, we propose a model in which PDA NPs enter the endosome–lysosome system via endocytosis and act as a mild exogenous stimulus that engages compensatory metabolic and secretory programs within lysosomes.

## 5. Conclusion

In summary, PDA participates throughout the entire hPSC-ALOs differentiation process by leveraging its tunable physical forms and intrinsic pro-adhesive and antioxidant properties, thereby systematically optimizing differentiation efficiency and functional maturation. Nevertheless, the current differentiation framework still relies on animal-derived matrices such as Matrigel, which limits translational applicability. Future efforts may therefore focus on using PDA coatings or PDA NPs as modular intermediates to engineer xeno-free cell culture substrates and fully defined 3D matrices. Such PDA-enabled, animal-free culture systems would represent an important step toward the standardization and clinical translation of hPSC-derived ALOs.

## Acknowledgements

This study was supported by grants from the National Key Research and Development Program of China (2021YFA1101304), Pearl River Talent Recruitment Program (2021QN02Y572 to H.M.), Guangzhou National Laboratory-State Key Laboratory of Respiratory Disease (GZNL-SKLRD) Joint Funding Program (GZNL2025B01010 to H.L.), Major Project of Guangzhou National Laboratory (SRPG22-021, GZNL2025C03007 and GZNL2024A03012 to H.L.), General Program of the National Natural Science Foundation of China (NSFC) (32570979 to H.L.).

## Author Contributions

Conceptualization: H.M. and H.L.; Methodology: R.L., Y.C. and H.M.; Investigation: R.L., Y.C., Z.L., H.Z. and J.Y.; Validation: Y.C., Z.L., C.Z. and H.Z.; Formal analysis: R.L., Y.C. and H.M.; Data curation: R.L., Y.C. and C.Z.; Visualization: R.L.and H.M.; Resources: H.L., T.X. and C.D.; Supervision: H.L. and T.X.; Project administration: H.M. and H.L.; Funding acquisition: H.M., H.L. and T.X.; Writing - original draft: R.L. and H.M.; Writing - review & editing: H.M. and H.L.. All authors reviewed and approved the final version of the manuscript.

## Data availability

Data will be made available on request.

