## Supplementary Figures 1 to 3; Supplementary Tables 1 to 4 for "Stage-adaptive integration of polydopamine promotes human pluripotent stem cell-derived alveolar organoids differentiation and maturation"

This file includes:

Supplementary Figures 1 to 3

Supplementary Tables 1 to 4

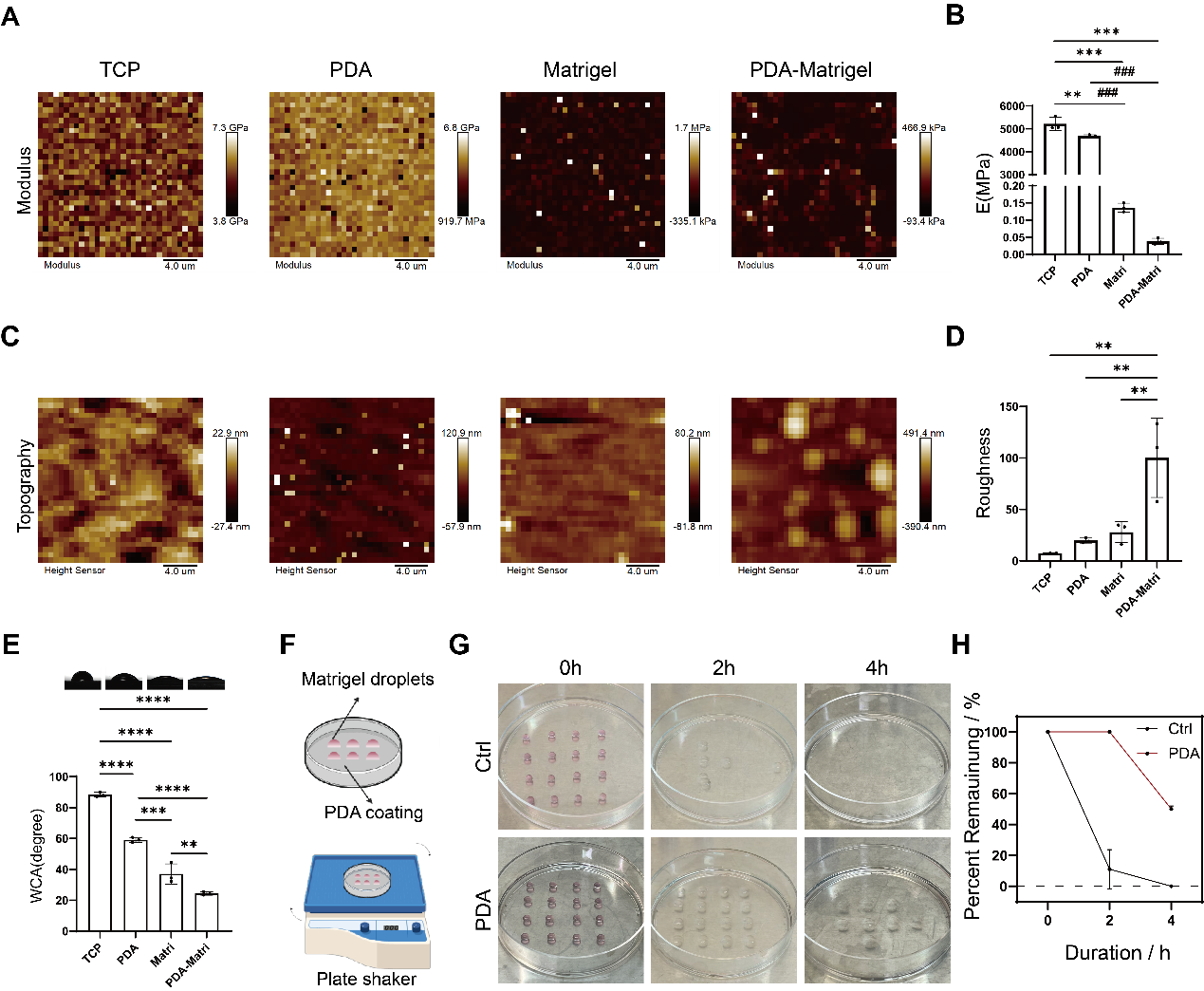

Fig.S1 Physiochemical characterization of PDA coating. A) Representative AFM maps displaying the spatial distribution of the Young’s modulus (E) on tissue culture plate (TCP), PDA-coated TCP (PDA), Matrigel-coated TCP (Matrigel), and PDA-Matrigel-coated TCP (PDA-Matrigel). Scale bar, 4 µm; B) Quantification of AFM-derived Young’s modulus for the indicated substrates; C) Representative AFM topography (height) of TCP, PDA, Matrigel and PDA-Matrigel. Scale bar, 4 µm; D) Quantification of surface roughness derived from AFM topography; E) Water contact angle (WCA) measurements evaluating surface wettability of TCP, PDA, Matrigel and PDA-Matrigel; F) Schematic illustration of the Matrigel droplet adhesion assay: an array of Matrigel droplets was deposited onto Ctrl or PDA-coated plates and subjected to orbital shaking; G) Representative images of Matrigel droplets retention on Ctrl and PDA substrates after agitation for 0, 2, and 4 h; H) Quantification of the percentage of Matrigel droplets remaining over time during agitation. Statistical significance is indicated as shown (**P < 0.01, *** and ### P < 0.001; # indicates comparisons as labeled).

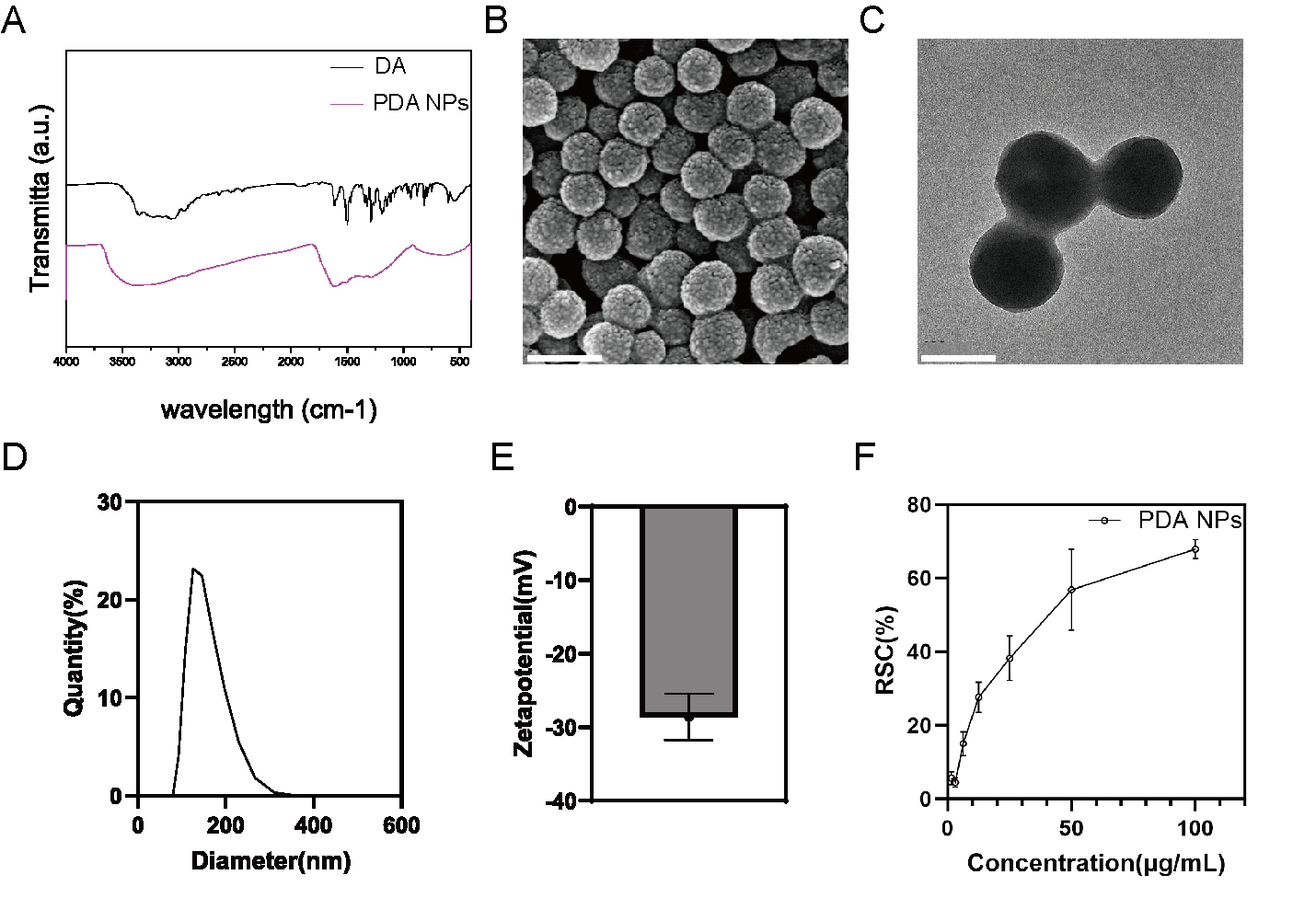

Fig.S2 PDA nanoparticles characteristics. A) Fourier-transform infrared (FTIR) spectra of dopamine and PDA NPs; B) Representative SEM image of PDA NPs, scale bar: 200 nm; C) Representative TEM image of PDA NPs, scale bar: 100nm; D) DLS analysis of particle size distribution of PDA NPs. E) Zeta potential of PDA NPs measured in aqueous suspension. F) DPPH radical scavenging capacity (RSC) assay of PDA NPs at different concentrations.

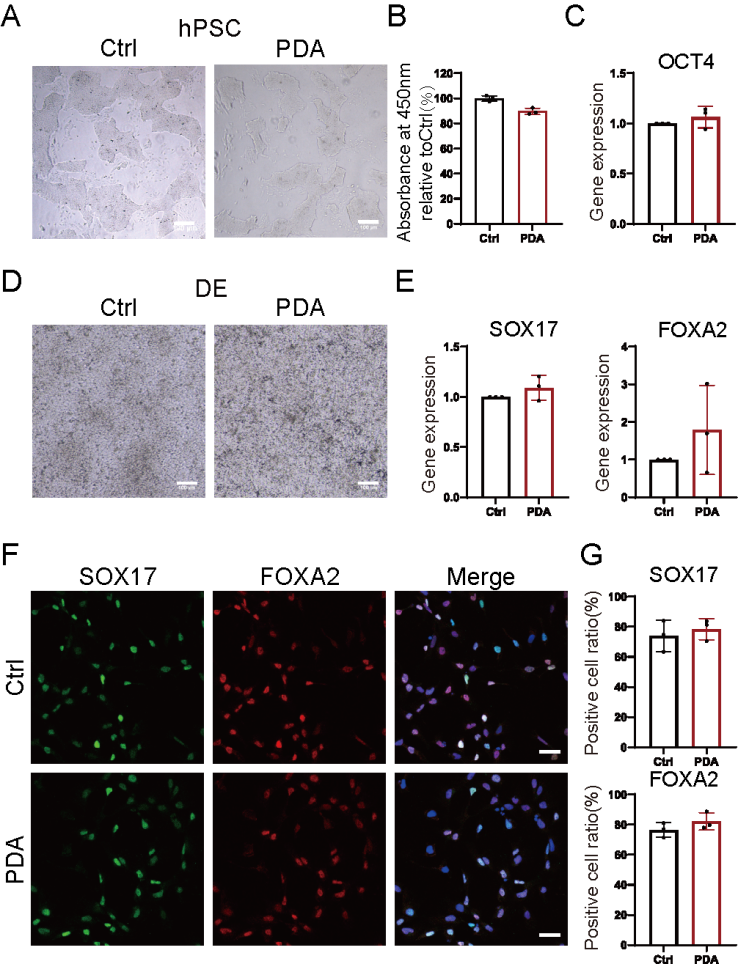

Fig.S3 PDA coating maintains hPSC proliferation and DE differentiation. A) Representative bright-field images of hPSCs cultured on Ctrl and PDA-coated substrates, scale bar: 100µm; B) CCK-8 assay of hPSC viability on Ctrl versus PDA-coated substrates; C) qPCR analysis of the OCT4 expression of hPSC cultured on Ctrl and PDA-coated substrates; D) Representative bright-field images of DE cells differentiated on Ctrl or PDA-coated substrates, scale bar: 100µm; E) qPCR analysis of DE marker genes SOX17 and FOXA2 expression in DE cells generated on Ctrl or PDA-coated substrates; F) Immunofluorescence imaging of SOX17 (green) and FOXA2 (red), (nuclei: blue. scale bar: 50µm); G) Quantification of SOX17 positive and FOXA2 positive cell percentages from (F).

**Supplementary Table 1. Reagents used in culture media**

| **Reagent name** | **Vandor (Cat#)** |
| --- | --- |
| MCDB131 | Thermo Fisher (Cat#10372019) |
| Advanced DMEM/F12 | Thermo Fisher (Cat#12634010) |
| DMEM/F12 medium | Thermo Fisher (Cat#10565018) |
| Glucose | Sigma (Cat#G7528-1kg) |
| Sodium bicarbonate (NaHCO_3_) | Sigma (Cat#6014-500g) |
| Bovine serum albumin (BSA) | Proliant (Cat#69700) |
| Penicillin/Streptomycin (P/S) | Thermo Fisher (Cat#15140122) |
| GlutaMAX | Thermo Fisher (Cat#35050061) |
| B27 | Thermo Fisher (Cat#17504044) |
| N-2 | Thermo Fisher (Cat#17502048) |
| ITS-X | Thermo Fisher (Cat#51500056) |
| HEPES | Thermo Fisher (Cat#15630080) |
| Activin A | StemCell (Cat#78001) |
| CHIR-99021 (CHIR) | MCE (Cat#HY-10182) |
| SAG | MCE (Cat#HY-12848) |
| SB-431542 (SB) | MCE (Cat#HY-10431) |
| Noggin | MCE (Cat#HY-P7051A) |
| FGF4 | MCE (Cat#HY-P7014) |
| DAPT | MCE (Cat#HY-13027) |
| BMP4 | MCE (Cat#HY-P7007) |
| FGF7 | MCE (Cat#HY-P70597) |
| FGF10 | MCE (Cat#HY-P70695) |
| Retinoic acid (RA) | Sigma (Cat#R2625) |
| Dexamethasone (DXMS) | MCE (Cat#HY-14648) |
| 8-Br-cAMP (8-Br) | MCE (Cat#HY-12318) |
| 3-isobutyl-1-methylxanthine (IBMX) | MCE (Cat#HY-12318) |

**Supplementary Table 2. Detailed medium formulation of hPSC-derived ALOs**

| Media | DE  Day 1 | DE  Day 2 | DE  Day 3 | AFE  Day 4-6 | LPC  Day 7-14 | ALO  Day 15-28 |
| --- | --- | --- | --- | --- | --- | --- |
| Basel media | MCDB131  Glucose ( 1.8 mg/mL)  NaHCO_3_ (1.5 mg/mL)  BSA (5 mg/mL)  GlutaMAX (1%)  P/S (1%) | | | Advanced DMEM/F12  B27 (2%)  N-2 (1%)  HEPES (10 mM)  GlutaMAX (1%)  P/S (1%) | | DMEM/F12  B27 (1%)  BSA (0.25%)  P/S (1%)  ITS-X (50 nM) |
| Add on day of use | 100 ng/mL Activin A | | | 10 μM SB  200 ng/mL Noggin  1 μM SAG  500 ng/mL FGF4  2 μM CHIR | 3 μM CHIR  10 ng/mL FGF7  10 ng/mL FGF10  20 μM DAPT  20 ng/mL BMP4  0.5 μM RA | 50nM DXMS  100nM 8-Br  100nM IBMX  3μM CHIR  10μM SB  10 ng/mL FGF7 |
|  | 3μM Chir | 0.1μM Chir |  |  |  |  |

**Supplementary Table 3. Primers used in the present study**

| **Gene** | **Forward (5’-3’)** | **Reverse (5’-3’)** |
| --- | --- | --- |
| *OCT4* | CCGAAAGAGAAAGCGAACCAG | ATGTGGCTGATCTGCTGCAGT |
| *SOX17* | GCATGACTCCGGTGTGAATCT | TCACACGTCAGGATAGTTGCAGT |
| *FOXA2* | GGGAGCGGTGAAGATGGA | TCATGTTGCTCACGGAGGAGTA |
| *NKX2-1* | CGGCATGAACATGAGCGGCAT | GCCGACAGGTACTTCTGTTGCTTG |
| *SOX2* | GCTTAGCCTCGTCGATGAAC | AACCCCAAGATGCACAACTC |
| *SOX9* | GACTACACCGACCACCAGAACTCC | CTGAGCTCGGCGTTGTG |
| *CDX2* | GGGCTCTCTGAGAGGCAGGT | GGTGACGGTGGGGTTTAGCA |
| *PAX6* | CGAATTCTGCAGGTGTCCAA | ACAGACCCCCTCGGACAGTAAT |
| *PAX8* | TGCCTCACAACTCCATCAGA | CAGGTCTACGATGCGCTG |
| *SFTPB* | TGCCTGGACCACCTCATCCTTG | GTCCTCACACTCTTGGCATAGG |
| *SFTPC* | AGCAAAGAGGTCCTGATGGA | CGATAAGAAGGCGTTTCAGG |
| *HOPX* | GCCTTTCCGAGGAGGAGAC | TCTGTGACGGATCTGCACTC |
| *AGER* | GCCACTGGTGCTGAAGTGTA | TGGTCTCCTTTCCATTCCTG |
| *GAPDH* | TGCACCACCAACTGCTTAGC | GGCATGGACTGTGGTCATGAG |

**Supplementary Table 4. Primary and secondary antibodies used in the present study**

| **Protein Name** | **Antibody** | **Vendor(cat#)** | **Application** |
| --- | --- | --- | --- |
| SOX17 | Anti-SOX17 antibody [OTI3B10] | Abcam (Ab84990) | IF (1:200) |
| FOXA2 | Anti-FOXA2 antibody [EPR4466] | Abcam (Ab108422) | IF (1:400) |
| NKX2-1 | Anti-TTF1/Nkx2-1 antibody [EP1584Y] | Abcam (Ab76013) | IF (1:250),  WB (1:2000) |
| SOX9 | Anti-SOX9 antibody [3C10] - BSA and Azide free | Abcam (Ab76997) | IF (1:500),  WB (1:2000) |
| SOX2 | Anti-SOX2 antibody | Abcam (Ab239218) | IF (1:50),  WB (1:2000) |
| AGER | Human/Mouse/Rat RAGE/AGER Antibody | R&D Systems (AF1145) | IF (1:250) |
| SFTPB | SP-B Antibody (F-2) | Santa Cruz Biotechnology (Sc-133143) | IF (1:200),  WB (1:1000) |
| ABCA3 | Anti-ATP-binding cassette sub-family A member 3 antibody | Abcam(ab99856) | IF (1:200) |
| β-tubulin | β-Tubulin Mouse mAb (HRP Conjugated) | Beyotime (AF2839) | WB (1:5000) |
| β-actin | Beta Actin Monoclonal antibody | proteintech(66009-1-Ig) | WB (1:5000) |
| Secondary antibodies | Donkey anti-Rabbit IgG (H+L) Highly Cross-Adsorbed Secondary Antibody, Alexa Fluor 488 | Invitrogen (A-21206) | IF (1:500) |
|  | Donkey anti-Mouse IgG (H+L) Highly Cross-Adsorbed Secondary Antibody, Alexa Fluor 488 | Invitrogen (A-21202) | IF (1:500) |
|  | Donkey anti-Rabbit IgG (H+L) Highly Cross-Adsorbed Secondary Antibody, Alexa Fluor546 | Invitrogen (A-10040) | IF (1:500) |
|  | Donkey anti-Goat IgG (H+L) Cross-Adsorbed Secondary Antibody, Alexa Fluor 546 | Invitrogen (A-11056) | IF (1:500) |
|  | Donkey anti-Mouse IgG (H+L) Highly Cross-Adsorbed Secondary Antibody, Alexa Fluor 647 | Invitrogen (A-31571) | IF (1:500) |
|  | HRP-Donkey Anti-Goat IgG (H+L) | proteintech (SA00001-3) | WB (1:5000) |
|  | HRP-conjugated Affinipure Goat Anti-Mouse IgG (H+L) | proteintech (SA00001-1) | WB (1:5000) |
|  | HRP-conjugated Affinipure Goat Anti-Rabbit IgG (H+L) | proteintech (SA00001-2) | WB (1:5000) |
